# Essential Role for Trf2 in Cardiac Development and Function

**DOI:** 10.1101/2025.09.24.677933

**Authors:** Ali Hakim Shoushtari, Aysenur Danis, Michael Wayne Stoner, Jeffrey Baust, Andrea Sebastiani, Paramesha Bugga, Janet R Manning, Bellina A S Mushala, Nisha Bhattarai, Imad Al Ghouleh, Iain Scott, Maryam Sharifi-Sanjani

## Abstract

Telomere Repeat-binding Factor 2 (Trf2) is essential for protecting our telomeres. While Trf2 global deletion is lethal, its role in organ-specific development, particularly in the heart, remains less understood. In this study, we investigated the role of Trf2 in cardiac development and function. Our studies reveal that cardiomyocyte (CM)-specific loss of Trf2 leads to profound defects in heart morphology, including impaired ventricular wall formation and compromised CM proliferation, concurrent with no CM telomere length attrition. Further, *in vivo* functional assessment and molecular analyses of CM-Trf2 deficient ventricles revealed severe cardiac dysfunction and, interestingly, altered nuclear envelope gene expression, respectively. Our work provides new insights into the essential role of Trf2 in heart development and function, and potential avenues for therapeutic intervention targeting telomere biology.

---

Telomere Repeat-binding Factor 2 (Trf2) is a critical member of the shelterin complex that protects telomeres^1^. While Trf2 global deletion is lethal, its role in organ-specific development, particularly in the heart, remains unexplored. Recent research has revealed new roles for Trf2 that vary across tissues; in non-cardiomyocyte (CM) cells, Trf2 has been implicated in DNA damage responses^1^, transcriptional regulation^2^, interaction with the nuclear lamina protein lamin A/C^1^, and the regulation of cell morphology and death in iPSCs from Duchenne Muscular Dystrophy patients^3^. Here, we show that CM-specific Trf2 deficiency induces profound developmental heart defects and compromises cardiac function; effects not attributable to Trf2’s canonical role in telomere length regulation. These findings reveal a critical, previously unrecognized role for Trf2 in supporting cardiac cellular and functional integrity.

To assess the role of Trf2 in the heart, we generated CM-specific Trf2 deficient mice. Animal experiments were approved by the University of Pittsburgh Institutional Animal Care and Use Committee. Trf2 floxed mice (Trf2^flx/flx^) (B6;129P2-Terf2tm1Tdl/J) were crossed with α-myosin-heavy chain (α-MyHC)-Cre mice (B6.FVB-Tg(Myh6-cre)2182Mds/J) (The Jackson Laboratory), followed by breeding hemizygous and homozygous α-MyHC-Cre;Trf2^flx/-^ with homozygous Trf2^flx/flx^ mice to generate male and female CM-specific hemizygous cTrf2^flx/-^ and homozygous cTrf2^flx/flx^ mice. Of 198 pups, no CM-specific Trf2-null (cTrf2^flx/flx^) mice were obtained, indicating complete embryonic lethality. Immunofluorescent staining of cTrf2^flx/flx^ embryonic hearts at day 14.5 (E14.5) revealed pronounced structural abnormalities including thinned ventricular walls, and a marked reduction in proliferating CMs evidenced by a ∼50% reduction in Ki67^+^ CMs (Figure 1A). These observations suggest impaired myocardial compaction and probable pre-birth cardiac rupture, demonstrating the absolute requirement of embryonic Trf2 for survival and cardiac morphogenesis.

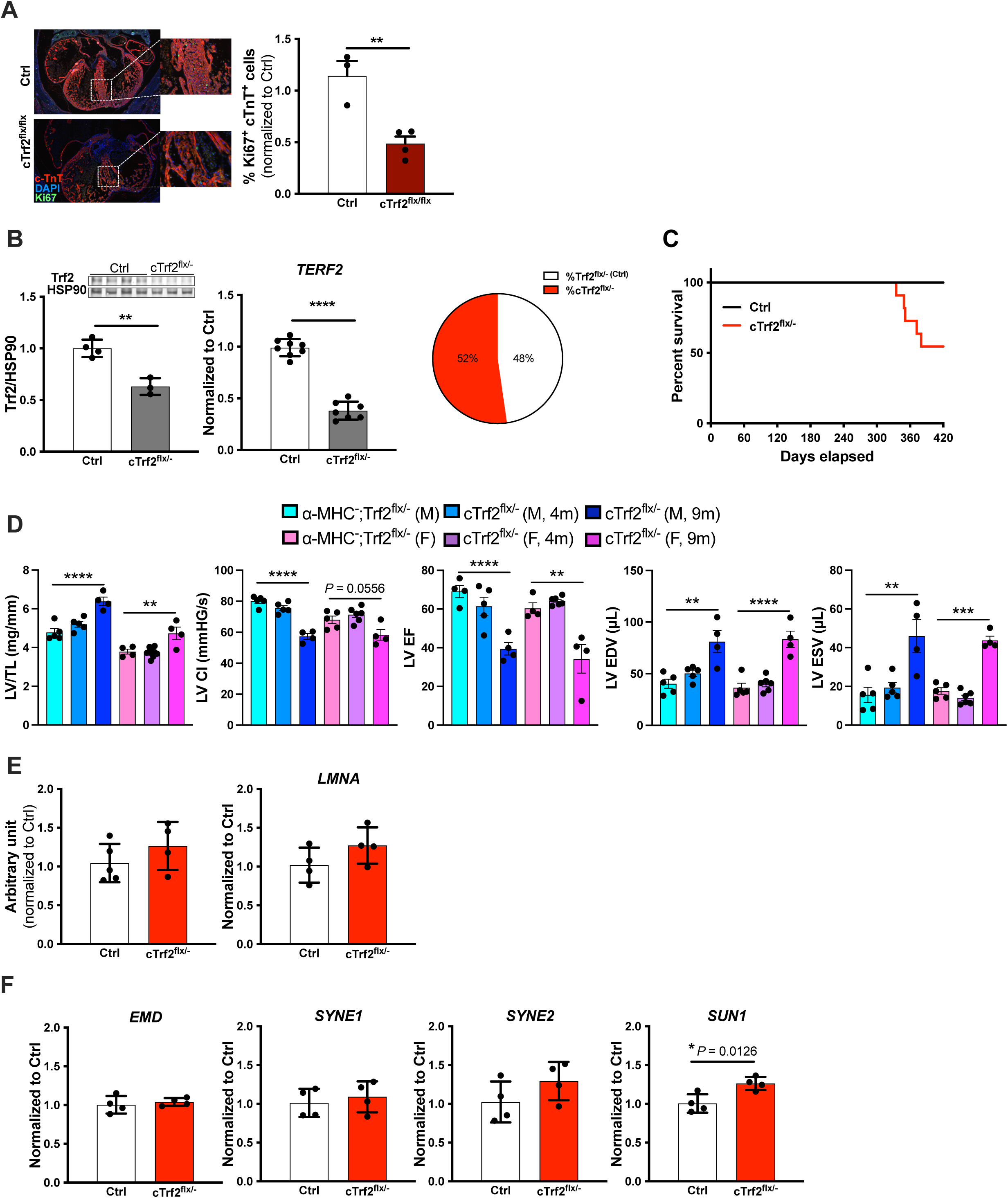
(**A**) Representative immunofluorescence images and quantification of E14.5 littermate control (Ctrl; α-MHC^**-**^ ;Trf2^flx/-^) and cTrf2^flx/-^ mice hearts; hearts stained for cardiac troponin T (cTnT, red), Ki67 (green), and DAPI (blue); n = 3-4 (**B**) Trf2 protein and mRNA (TERF2) levels in 4m Ctrl and cTrf2^flx/-^ mice LVs and pups genotype percentages (%) from 88 born (pie plot); n=3-8 (**C**) Kaplan-Meier survival curve (with 95% CIs) of Ctrl and cTrf2^flx/-^ mice showing marked change in survival in cTrf2^flx/-^ mice occurring around week 47; n = 15-17 (**D**) LV weight/tibia length (LV/TL) and *in vivo* cardiac hemodynamic (pressure-volume loop) assessment showing LV contractile index (CI), ejection fraction (EF), end-diastolic volume (EDV), and end-systolic volume (ESV) in M (blue graphs) and F (pink graphs) Ctrl and cTrf2^flx/-^ mice at 4m and 9m of age; n = 4-6 (**E**) CM telomere length quantification of Ctrl and cTrf2^flx/-^ E14.5 hearts and lamin A/C transcript levels in Ctrl and cTrf2^flx/-^ LVs; n = 4-5 (**F**) Transcript levels of emerin (EMD), Nesprin-1 (Syne1), Nesprin-2 (Syne2), and SUN1 in Ctrl and cTrf2^flx/-^ LVs; n = 4. Experiments performed in male (M) and female (F) mice; Data were collected with blinding and are expressed as mean ± SEM. Statistical significance was determined by one-way ANOVA with post-hoc Bonferroni test, or t-test, using GraphPad Prism; *P* < 0.05 was considered significant. Significance: \**P* < 0.05, ** *P* <0.005, *** *P* <0.0005, **** *P* < 0.0001.

In contrast, hemizygous cTrf2^flx/-^ mice were viable, with comparable birth percentages to their corresponding control littermates (Figure 1B). However, cTrf2^flx/-^ mice exhibited premature death (Figure 1C), left ventricular hypertrophy, dilated ventricles, and impaired contractile function at nine months (Fig. 1D). As emerging studies indicate diverse functions for Trf2 across various tissues, we aimed to understand the mechanism(s) through which Trf2 exerts its effect in the heart. Telomere shortening primarily occurs in proliferative cells, and since CMs proliferate mainly during embryogenesis (with proliferation declining shortly after birth), we assessed CM telomere length at E14.5 using immunofluorescent staining and fluorescence in situ hybridization. Interestingly, we found that CM-Trf2 deficiency did not induce telomere shortening at this key CM proliferation stage (Figure 1E). This finding is consistent with our report^4^ showing the absence of CM telomere shortening in certain human cardiomyopathies in which Trf2 expression is decreased. Furthermore, given that Trf2 interacts with lamin A/C (a protein linked to laminopathies and dilated cardiomyopathy; DCM), and that disruptions in this interaction influence telomere length, we examined lamin A/C expression but found no change (Figure 1E). As previously reported, global Trf2 deletion is embryonically lethal; this lethality may reflect the embryonic effect of Trf2 on CM proliferation, or telomere length-related consequences in other cell types. We also must consider that in the case of our studies, residual Trf2 in cTrf2^flx/-^ mice may be adequate to maintain telomere length, whereas complete CM-Trf2 deletion could trigger telomere attrition. However, the viability of global Telomerase Reverse Transcriptase (Tert; the enzyme responsible for maintaining telomere length) knockout mice argues against telomere attrition as the sole explanation.

To look beyond previously reported Trf2 mechanisms and its telomere-related role, we examined the expression of emerin (EMD), a lamin A/C interacting protein that contributes to DCM, but found no change (Figure 1F). We then examined the expression of two lamin A/C- and emerin-interacting nuclear membrane proteins, Nesprin-1 (SYNE1) and Nesprin-2 (SYNE2), that are also linked to DCM. Although we did not find any changes in SYNE1 and SYNE2 expression, we found significantly increased levels of the nuclear lamina-interacting inner nuclear envelope protein Sun1 (Figure 1F). Importantly, Sun1 is a core component of the LINC complex, which physically connects the nuclear envelope to the cytoskeleton, mediating mechanical and force transmission. Sun1 depletion in lamin A-related DCM is reported to enhance longevity and ventricular function^5^, both altered parameters in our Trf2-deficient mice. Our finding suggests that elevated Sun1 expression may underlie the observed functional loss and ventricular dilation in Trf2-deficient hearts, and implies a regulatory role for Trf2 in CM gene expression. Further investigation is warranted to elucidate the spectrum of Trf2’s gene targets, and clarify whether these transcriptional effects drive, or result from, cardiac remodeling.

In conclusion, we establish Trf2 as a pivotal regulator of cardiac development and function via CM gene regulation. Loss of Trf2 in embryonic CMs causes profound structural defects driven by impaired CM proliferation, while Trf2-deficient adults exhibit premature mortality and severe systolic and diastolic dysfunction. These findings reveal a previously unrecognized role for Trf2 beyond telomere protection, implicating nuclear envelope gene dysregulation in cardiac pathology, and underscoring the diverse context- and cell type-specific functions of telomeric proteins.

## Acknowledgement

We thank the Department of Cell biology, Center for Biologic Imaging, University of Pittsburgh for providing advanced imaging facility access and their help with imaging experiments.

